# Conformational dynamics of the HIV-1 envelope glycoprotein from CRF01_AE is associated with susceptibility to antibody-dependent cellular cytotoxicity

**DOI:** 10.1101/2024.08.22.609179

**Authors:** Marco A. Díaz-Salinas, Debashree Chatterjee, Manon Nayrac, Halima Medjahed, Jérémie Prévost, Marzena Pazgier, Andrés Finzi, James B. Munro

## Abstract

The HIV-1 envelope glycoprotein (Env) is expressed at the surface of infected cells and as such can be targeted by non-neutralizing antibodies (nnAbs) that mediate antibody-dependent cellular cytotoxicity (ADCC). Previous single-molecule Förster resonance energy transfer (smFRET) studies demonstrated that Env from clinical isolates predominantly adopt a “closed” conformation (State 1), which is resistant to nnAbs. After interacting with the cellular receptor CD4, the conformational equilibrium of Env shifts toward States 2 and 3, exposing the coreceptor binding site (CoRBS) and permitting binding of antibodies targeting this site. We showed that the binding of anti-CoRBS Abs enables the engagement of other nnAbs that target the cluster A epitopes on Env. Anti-cluster A nnAbs stabilize an asymmetric Env conformation, State 2A, and have potent ADCC activity. CRF01_AE strains were suggested to be intrinsically susceptible to ADCC mediated by nnAbs. This may be due to the presence of a histidine at position 375, known to shift Env towards more “open” conformations. In this work, through adaptation of an established smFRET imaging approach, we report that the conformational dynamics of native, unliganded HIV-1_CRF01_AE_ Env indicates frequent sampling of the State 2A conformation. This is in striking contrast with Envs from clades A and B, for example HIV-1_JR-FL_, which do not transition to State 2A in the absence of ligands. These findings inform on the conformational dynamics of HIV-1_CRF01_AE_ Env, which are relevant for structure-based design of both synthetic inhibitors of receptor binding, and enhancers of ADCC as therapeutic alternatives.

**IMPORTANCE:** A concerning increase in infections with HIV-1_CRF01_AE_ has occurred globally and regionally in recent years, especially in Southeast Asia. Despite the advances made in understanding HIV-1 Env conformational dynamics, the knowledge about Env from HIV- 1_CRF01_AE_ is limited. Here, we demonstrate that HIV-1_CRF01_AE_ Env readily samples an open conformation (State 2A), which is susceptible to ADCC. This is in contrast with the subtypes previously studied from HIV-1 group M that rely on anti-cluster A antibodies to adopt State 2A. These findings are relevant for the structure-based design of novel synthetic inhibitors of CD4 binding and enhancers of ADCC for elimination of infected cells.

## INTRODUCTION

The RV144 HIV-1 vaccine trial in Thailand, which concluded in 2009, elicited a 31.2% protective efficacy. Subsequent analyses indicated that this modest protection was correlated with antibodies (Abs) with Ab-dependent cellular cytotoxicity (ADCC) activity specific to the HIV-1 envelope glycoprotein (Env) in a subset of individuals with low plasma IgA (1, 2). This suggests that ADCC may have contributed to the protection observed in the RV144 trial. HIV-1 strains of the circulating recombinant form AE (CRF01_AE) predominate the AIDS epidemic in Southeast Asia (3). Therefore, the RV144 trial utilized glycoproteins from two CRF01_AE strains as immunogens.

Moreover, the prevalence of HIV-1 CRFs has risen in recent years, most significantly in Southeast Asia (4). For these reasons, detailed investigation of Env from HIV-1 CRFs is warranted. While advances in the understanding of Env conformational dynamics have been achieved using virological and biophysical approaches, these studies have focused on HIV-1 subtypes A and B (5–11). A similar elucidation of the dynamics of Env from HIV-1 CRFs has not been reported. However, prior studies demonstrated an inherent susceptibility of HIV-1_CRF01_AE_ to ADCC, which begins to explain the results of the RV144 trial (12). Subsequent structural investigation of HIV-1_CRF01_AE_ Env indicated features that are distinct from other subtypes and perhaps enable conformations related to recognition by Abs with ADCC activity (13). In the present study, we explore the conformational features of Env from HIV-1_CRF01_AE_ and their relationship to ADCC mediated by plasma from people living with HIV (PLWH).

The first step in replication of HIV-1 is the binding of Env to the cellular receptor CD4. Env is synthesized as the gp160 precursor, which is trimerized and glycosylated in the endoplasmic reticulum of infected cells (14, 15), followed by proteolytic processing by host furin-like proteases in the Golgi apparatus (16–18). The resulting cleaved and mature Env trimer is comprised of three gp120 subunits, which are non-covalently associated with three transmembrane gp41 subunits [(gp120-gp41)_3_] (19–21). Mature Env is present on virions as well as exposed on the surface of infected cells, making it the primary target of host Abs. Some Abs neutralize the virus (NAbs) by blocking Env’s interaction with receptors or inhibiting conformational changes needed to promote fusion of the viral and cellular membranes. Other Abs that are frequently elicited during HIV-1 infection, including in people leaving with HIV (PLWH), are non-neutralizing (nnAbs) since they recognize Env targets occluded within closed Env conformations. Certain classes of nnAbs, however, can induce the death of infected cells through ADCC, provided Env samples an “open” conformation (22).

Single-molecule Förster resonance energy transfer (smFRET) imaging studies demonstrated that Env is highly dynamic, transitioning from a “closed” conformation (State 1) to an “open” conformation (State 3), which is promoted through the interaction with CD4. An asymmetric intermediate (State 2) of Env can be observed during the transition from State 1 to State 3 (9, 11). The Env conformational equilibrium from primary HIV-1 isolates of clades A and B favor State 1 in the absence of ligands, which confers resistance to most Abs, especially those that target CD4-induced (CD4i) epitopes (11, 23). Nonetheless, some broadly neutralizing Abs (bNAbs) preferentially bind this closed conformation (7, 11). However, after interacting with cellular CD4, Env adopts State 3, exposing cryptic epitopes including the coreceptor-binding site (CoRBS) and cluster A region, which can be targeted by nnAbs to promote ADCC (5, 23–28).

CD4-mimetic compounds (CD4mcs) are small molecules designed to target specifically the CD4 binding cavity within HIV-1 Env. CD4mcs can induce conformational changes in Env that sensitize it to recognition by nnAbs (25, 26). In the presence of soluble CD4 (sCD4) or CD4mcs, anti-CoRBS and anti-cluster A Abs stabilize State 2A, which is an asymmetric Env conformation associated with increased ADCC responses *in vitro* and Fc-effector functions *in vivo* (8, 25, 29–31).

The findings presented here indicate that native HIV-1_CRF01_AE_ Env intrinsically presents the State 2A conformation, which is susceptible to ADCC even in the absence of CD4 or CD4mcs. This contrasts with clade-B HIV-1_JR-FL_ Env, which depends on incubation with CD4 or CD4mcs, and antibodies targeting the CoRBS to adopt State 2A (8, 25). Interaction of HIV-1_CRF01_AE_ Env with CD4 and CoRBS Abs further stabilized State 2A. The conformational features of HIV-1_CRF01_AE_ Env warrants further research to identify the structural determinants or elements that govern its dynamic equilibrium.

Targeting cells infected with HIV-1_CRF01_AE_ could represent a promising strategy for elimination of infected cells (31–33).

## RESULTS

### HIV-1_CRF01_AE_ is more susceptible to ADCC than a representative subtype B strain

We made a direct comparison of the susceptibility of infected cells to ADCC using representative infectious molecular clones (IMCs) from CRF01_AE (strain 703357) and subtype B (strain JR-FL). First, we evaluated the binding capacity of plasma from ten PLWH (Table 1). No significant differences between the two strains were observed (**Fig. 1A**). However, the ADCC responses to HIV-1_CRF01_AE_ were approximately two-fold higher than that observed with HIV-1_JR-FL_ strain (**Fig. 1B**). Because activation of the ADCC response has been associated with a specific conformation of HIV-1 Env that enables binding of a specific class of Abs, these results suggest that HIV-1_CRF01_AE_ Env may have distinct conformational features that confer the sensitivity to ADCC (5, 6).

**Fig 1.**
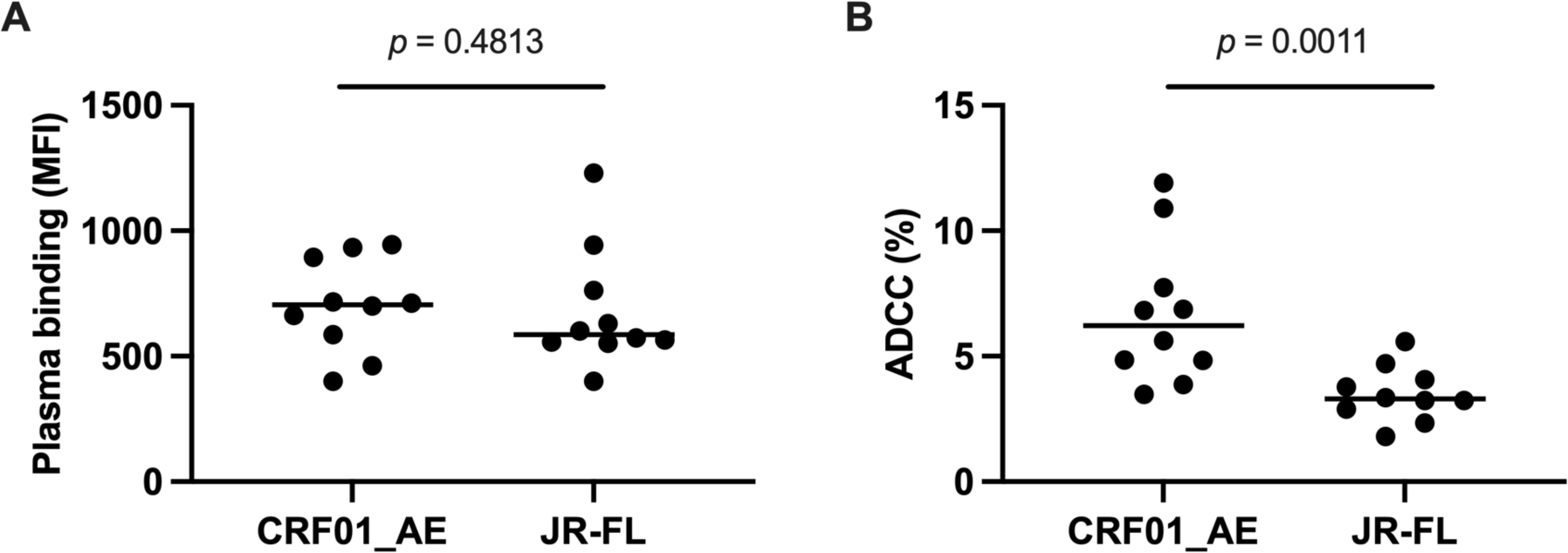
HIV-1_CRF01_AE_ strain 703357 is more susceptible to ADCC than HIV-1_JR-FL_. (**A**) Binding of plasma from PLWH to primary CD4+ T cells infected with the indicated HIV-1 strains was evaluated. Five independent experiments (n=5) were performed with each one of the ten plasma samples plotted as individual dots. Means are shown as horizontal bars. (**B**) ADCC responses to the indicated viral strains. Data are plotted as in (A). In this case, the number of independent experiments were 5 and 4 for HIV-1_CRF01_AE_ and HIV-1_JR-FL_, respectively. Statistical significance was determined through an unpaired two-tailed Mann-Whitney *t*-test and *p*-values < 0.05 were considered statistically significant.

**Table 1.** Characteristics of the cohort of people living with HIV, related to. **Fig. 1**.

| Donors | Sex | Age (years) | Days since infection | Days between inf. and ART | Viral load (copies/mL) | CD4 count (cells/mm<sup>3</sup>) |
| --- | --- | --- | --- | --- | --- | --- |
| P5 | M | 33 | 936 | 192 | 50 | 1149 |
| P8 | M | 42 | 999 | N/A | 28666 | 260 |
| P10 | M | 58 | 961 | 204 | 50 | 170 |
| P13 | M | 28 | 1194 | N/A | 809600 | 200 |
| P16 | F | 36 | 1576 | 833 | 50 | 570 |
| P25 | M | 33 | 1173 | N/A | 35922 | 421 |
| P30 | M | 40 | 1143 | N/A | 34759 | 691 |
| P43 | M | 34 | 856 | 1098 | 29234 | 410 |
| P44 | M | 34 | 1018 | 534 | 40 | 600 |
| P46 | M | 56 | 1005 | 986 | 40 | 780 |

### Modifications in HIV-1_CRF01_AE_ Env that enable site-specific fluorescent labeling do not affect viral infectivity

With the aim of visualizing the conformational dynamics of HIV-1_CRF01_AE_ Env, we adapted a previously validated smFRET imaging assay. Insertion of the A4 peptide (DSLDMLEW) and incorporation of non-natural amino acids (nnAAs) into HIV-1 Env facilitate fluorophore attachment. These methods have been applied with minimal effect on functionally to subtype-B HIV-1 strains NL4-3 and JR-FL, as well as the subtype-A strain BG505 (7–9, 11, 30). As for previous applications, we attached site-specifically fluorophores in the V1 and V4 loops of a single gp120 domain within HIV-1_CRF01_AE_ Env on the surface of pseudovirions (**Fig. 2A**). To this end, we inserted the A4 peptide next to V135 in V1 (V1-A4), which enabled enzymatic attachment of the LD650 fluorophore. We also substituted an amber stop codon for amino acid N398 in V4 of gp120 (V4- N398^TAG^). Suppression of the amber stop codon incorporates the nnAA TCO*, which facilitated Cy3 fluorophore attachment through copper-free click chemistry (**Fig. 2B**) (34).

**Fig 2.**
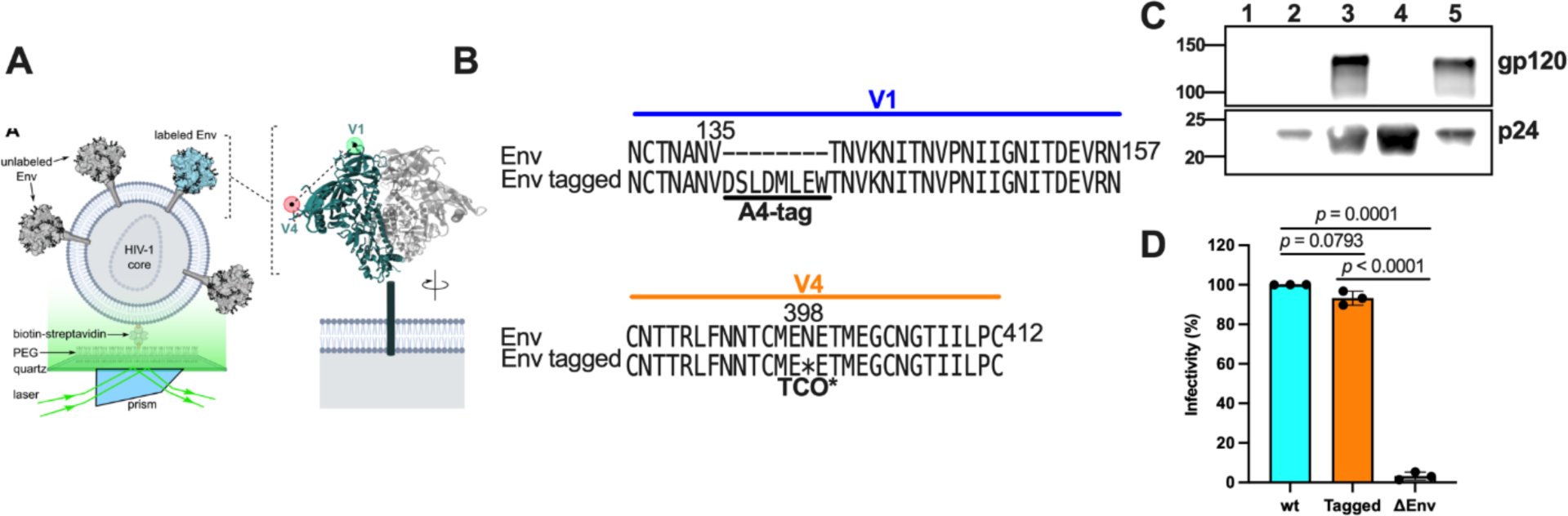
Engineering HIV-1_CRF01_AE_ Env for site-specific fluorescent labeling. (**A**) Schematic of the smFRET imaging assay. Pseudovirions with HIV-1_CRF01_AE_ Env (strain 92TH023) containing a single labeled gp120 domain were immobilized on quartz slides and imaged using TIRF microscopy. (**B**) Sequence alignments indicating sites of A4 peptide insertion into the V1 loop and TCO* substitution in the V4 loop for fluorophore attachment. (**C**) Qualitative detection of the indicated proteins from purified pseudovirions with HIV-1_CRF01_AE_ Env through immunoblots. Lane 1, mock-produced virus; lane 2, ι1Env virions; lane 3, wild-type Env pseudotyped virions; lane 4, Env V1- A4/V4-TAG (tagged) pseudotyped virions produced in the absence of the TCO* amino acid; lane 5, tagged Env pseudovirions produced in the presence of the TCO* amino acid. (**D**) Infectivity of lentiviruses with either wild-type HIV-1_CRF01_AE_ Env (wt), V1-A4/V4- TAG Env (tagged), or bald particles was evaluated in TZM-bI cells. Infectivity values are expressed as the percentage of wild-type Env and normalized to the expression level of gp120 and p24. Each point indicates the arithmetic mean of three technical replicates. Bars represent the average of three independent experiments per condition. Error bars reflect the standard error. The statistical significance was evaluated through parametric *t*-tests. *p*-values are indicated and those <0.05 were considered statistically significant.

We next confirmed full-length translation of the HIV-1_CRF01_AE_ Env containing the V1-A4 and V4-N398^TAG^ mutations (tagged) and its incorporation into virions. We evaluated through immunoblots the abundance of both full-length gp120 and the HIV-1 core capsid protein p24 in purified viral preparations (**Fig. 2C**). As expected, tagged gp120 was not detected in virions produced in the absence of the nnAA TCO* and the corresponding aminoacyl tRNA synthetase and suppressor tRNA, which codes for the amber stop codon. This indicates that readthrough of the amber codon in the V4 loop did not occur, resulting in the lack of Env incorporation into viral particles (**Fig. 2C**, top immunoblot, lane 4). However, in the presence of TCO*, the synthetase, and the suppressor tRNA, tagged gp120 was detected in virions at a comparable level as wild-type Env (**Fig. 2C**, top immunoblot, lane 5). We next verified that V1-A4/V4-N398^TAG^ modifications in Env do not alter virus infectivity. Virus preparations bearing wild-type or tagged Env showed no statistically significant difference in their infectivity in TZM-bI cells (**Fig. 2D**), suggesting that both incorporation of the A4 peptide in V1 and the nnAA TCO* in V4 does not affect the function of Env. Altogether, these data demonstrate that tagged Env is incorporated into pseudovirions and maintains native function during infection of cells.

### Native HIV-1_CRF01_AE_ Env intrinsically samples open conformations

We next sought to evaluate the conformational dynamics in real-time of individual HIV-1_CRF01_AE_ Env molecules on the surface of virions using smFRET imaging. To this end, we prepared virions bearing a single fluorescently labeled gp120 domain as described for Env from other HIV-1 strains (**Fig. 2A**) (7–9, 11). Labelled virions were immobilized on passivated quartz microscope slides and imaged using prism-based total internal reflection fluorescence (TIRF) microscopy. We used the well-characterized HIV-1_JR-FL_ Env as a point of comparison. As previously described, the application of hidden Markov modeling (HMM) for analysis of the smFRET trajectories enabled the identification of four FRET states (**Fig. 3A**). For both HIV-1_CRF01_AE_ and HIV-1_JR-FL_ Env, the predominant low-FRET value (0.22±0.1 FRET [mean ± standard deviation], State 1) is associated with a closed Env conformation (**Fig. 3A-B**, **Table 2**). Quantification of the mean occupancies in State 1 across the populations of molecules indicated 68±2% and 42±2% (*p* < 10^-4^) for HIV-1_JR-FL_ and HIV-1_CRF01_AE_, respectively. We also observed State 3 (0.45±0.1 FRET) for both strains, which is associated with an open Env conformation. We determined State 3 occupancies of 32±2% and 27±2% (*p* = 0.6) for HIV-1_JR-FL_ and HIV-1_CRF01_AE_, respectively. Consistent with previous reports, we detected minimal occupancy for HIV-1_JR-FL_ Env in States 2 and 2A (0.70±0.1 and 0.85±0.1 FRET, respectively). In striking contrast, HIV-1_CRF01_AE_ Env displayed 19±2% occupancy in State 2 and 12±1% in State 2A in the absence of bound ligands. These data demonstrate that HIV-1_CRF01_AE_ Env has greater intrinsic access to open conformations than HIV-1_JR-FL_ Env.

**Fig 3.**
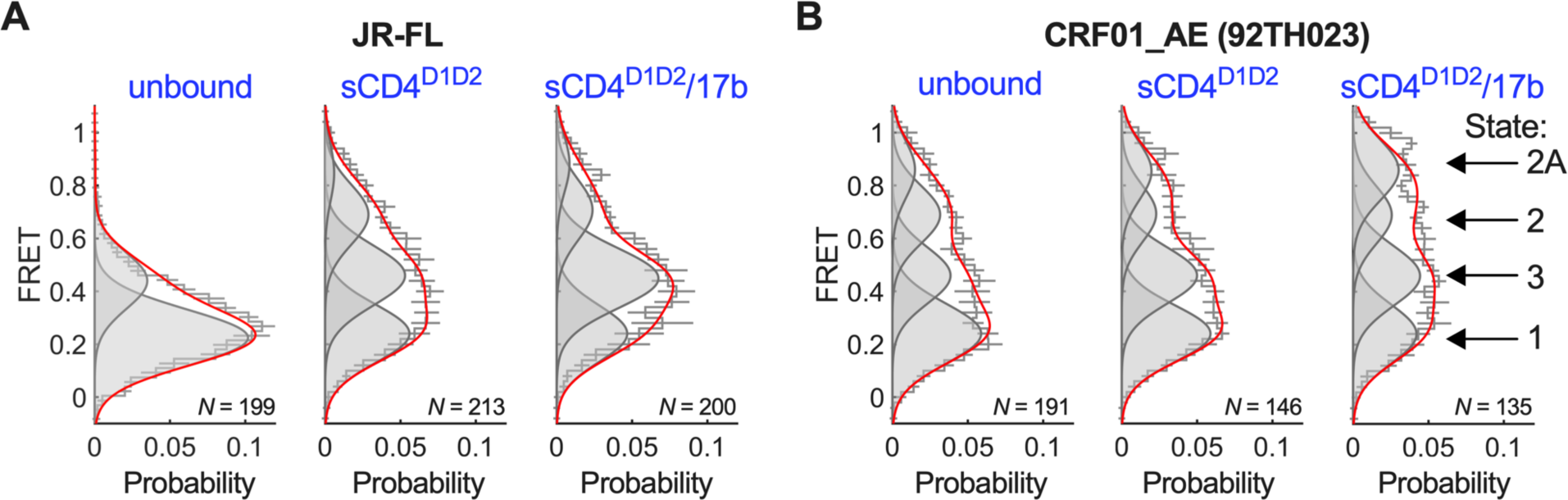
Conformational equilibrium of HIV-1_CRF01_AE_ Env. (**A**) FRET histograms from unbound HIV-1_JR-FL_ Env trimers, or Env pre-incubated with sCD4^D1D2^, or sCD4^D1D2^ and 17b, as indicated. FRET histograms are presented as the mean ± standard error determined from three technical replicates and the total number of smFRET traces used in the HMM analysis is shown (N). Overlaid on the histograms are four Gaussian distributions shown in grey and centered at 0.22 (State 1), 0.45 (State 3), 0.70 (State 2), and 0.85 (State 2A) FRET as determined through HMM analysis. The sum of the four Gaussians is shown in red. (**B**) The same data acquired for HIV-1_CRF01_AE_ Env trimers. Corresponding numeric FRET state occupancies are shown in **Table 2**.

**Table 2.** FRET-state occupancies for HIV-1 Env in the presence and absence of sCD4^D1D2^ and 17b.

| Strain/variant Env | Experimental condition | FRET state occupancies (%)<sup>a</sup> |  |  |  |
| --- | --- | --- | --- | --- | --- |
|  |  | State 1 | State 3 | State 2 | State 2A |
| JR-FL | Unbound | 68±2 | 32±2 | 0±2 | 0±1 |
|  | + sCD4<sup>D1D2</sup> | 35±2 | 44±2 | 18±2 | 3±1 |
|  | + sCD4<sup>D1D2</sup>/17b | 31±3 | 46±3 | 15±2 | 8±2 |
| CRF01_AE | Unbound | 42±2 | 27±2 | 19±2 | 12±1 |
|  | + sCD4<sup>D1D2</sup> | 39±2 | 33±2 | 18±2 | 10±2 |
|  | + sCD4<sup>D1D2</sup>/17b | 29±3 | 34±2 | 21±2 | 17±2 |

<sup>a</sup>Data are presented as mean ± standard error determined from the total population of traces analyzed.

We next asked if sCD4 consisting of soluble domain 1 and 2 (sCD4^D1D2^) or the anti-CoRBS 17b mAb, further stabilize open conformations. For both HIV-1_JR-FL_ and HIV- 1_CRF01_AE_ Env, the addition of sCD4^D1D2^ destabilized State 1 and promoted transition to the higher FRET states (**Fig. 3A-B**, **Table 2**). For HIV-1_JR-FL_ Env, we observed increased occupancy in States 2 and 3, as previously reported (11). sCD4^D1D2^ had only a modest effect on HIV-1_CRF01_AE_ Env conformation, with only a slight stabilization of State 3.

Addition of both sCD4^D1D2^ and 17b further promoted State 3 for HIV-1_JR-FL_ Env, as expected. In contrast, the predominant effect of sCD4^D1D2^/17b on HIV-1_CRF01_AE_ Env was to stabilize State 2A, increasing the occupancy to 17±2%. These data demonstrate that sCD4^D1D2^ only minimally promotes open conformations of HIV-1_CRF01_AE_ Env beyond that seen in the absence of ligands. However, HIV-1_CRF01_AE_ Env readily adopts State 2A in the presence of sCD4^D1D2^ and 17b. Access to State 2A correlates with the inherent sensitivity to ADCC seen for HIV-1_CRF01_AE_.

## DISCUSSION

During HIV-1 infection, the humoral response against Env mainly produces antibodies that are non-neutralizing. Despite the lack of neutralization, nnAbs can still trigger ADCC to clear infected cells, provided that Env is exposed in an “open” conformation (35). Env glycoproteins from most HIV-1 strains naturally adopt State 1, which is associated with a closed conformation (11), and confers resistance to nnAbs (23, 36). In contrast, previous functional studies suggested that Env glycoproteins from HIV-1_CRF01_AE_ subtypes intrinsically adopt open conformations even in the absence of CD4, CD4 mimetics, or anti-CoRBS mAbs (5, 6, 12). Recent insights from structural data further support this idea (13). Here, we have shown that plasma obtained from PLWH triggers ADCC against HIV-1_CRF01_AE_ infected cells to a greater extent than HIV- 1_JR-FL_ infected cells. We therefore sought to directly test the conformational equilibrium of HIV-1_CRF01_AE_ Env using smFRET imaging. We have demonstrated through real-time analysis of HIV-1_CRF01_AE_ Env conformational dynamics that this glycoprotein intrinsically samples open conformations in the absence of bound ligands. HIV-1_CRF01_AE 92TH023_ Env intrinsically adopts State 2A, which was previously linked to exposure of both the CoRBS and cluster A epitopes that are targeted by Abs with potent ADCC activity (8).

Addition of sCD4^D1D2^, with or without 17b, stabilized State 2A to a greater extent than seen for HIV-1_JR-FL_ Env. The results presented here were obtained with Env from HIV- 1_CRF01_AE_ strain 92TH023. Additional effort should be devoted to generalizing these results to Envs from additional CRF01_AE strains. Nevertheless, the data presented here provide a means of interpreting the inherent sensitivity of HIV-1_CRF01_AE_ to ADCC in terms of the conformation of Env. These data also provide new understanding for the role of vaccine-induced Abs that mediated ADCC during the RV144 trial in Thailand, where HIV-1_CRF01_AE_ predominates (5).To conclude, our data strongly underscore the importance of considering Env conformational diversity across different HIV-1 clades when designing more effective HIV-1 interventions and vaccine strategies. This is of particular importance for the development of tailored strategies for enhancing ADCC against HIV-1_CRF01_AE_, which offers promising avenues for the elimination of cells infected with this prevalent strain in Southeast Asia.

## MATERIALS AND METHODS

### Ethics statement

Written informed consent was obtained from all study participants, and the research adhered to the ethical guidelines of CRCHUM and was reviewed and approved by the CRCHUM Institutional Review Board (Ethics Committee approval number MP-02-2024-11734). The research adhered to the standards indicated by the Declaration of Helsinki. All participants were adults and provided informed written consent prior to enrollment, in accordance with the Institutional Review Board approval.

### Plasma samples

The FRQS-AIDS and Infectious Diseases Network supports a representative cohort of newly-HIV-infected subjects with clinical indication of primary infection [the Montreal Primary HIV Infection Cohort]. Plasma samples from ten deidentified PLWH donors were heat-inactivated and stored as previously described (24, 26).

### Cell lines and primary cells

ExpiCHO-S cells (Gibco, Thermo Fisher Scientific, Waltham, MA, USA) were cultured in ExpiCHO Expression media (Gibco, Thermo Fisher Scientific, Waltham, MA, USA) at 37 °C, 8% CO_2_ with orbital shaking according to manufacturer instructions. The cell line HEK293T-FIRB with enhanced furin expression was a kind gift from Dr. Theodore C. Pierson (Emerging Respiratory Virus section, Laboratory of Infectious Diseases, NIH, Bethesda, MD), and was cultured at 37°C, 5% CO_2_ in complete DMEM made of DMEM (Gibco, ThermoFisher Scientific, Waltham, MA, USA) supplemented with 10% (v/v) cosmic calf serum (Hyclone, Cytiva Life Sciences, Marlborough, MA, USA), 100 U/ml penicillin, 100 µg/ml streptomycin, and 1 mM glutamine (Gibco, ThermoFisher Scientific, Waltham, MA, USA) (37). The HeLa-derived TZM-bl cell line stably expressing high levels of CD4 and CCR5 receptors and bearing an integrated copy of the luciferase gene under the control of the HIV-1 long-terminal repeat was obtained from the former NIH AIDS Reagent Program (catalog ARP-8129) and cultured in the same conditions as HEK293T-FIRB cells (38).

Human peripheral blood mononuclear cells (PBMCs) from 3 HIV-negative individuals (3 males, age range 40-66 years) obtained by leukapheresis and Ficoll density gradient isolation were cryopreserved in liquid nitrogen until further use. Primary CD4+ T cells were purified from resting PBMCs by negative selection using immunomagnetic beads per the manufacturer’s instructions (StemCell Technologies, Vancouver, BC) and were activated with phytohemagglutinin-L (PHA-L, 10 μg/mL) for 48h and then maintained in RPMI 1640 (Thermo Fisher Scientific, Waltham, MA, USA) complete medium supplemented with 20% FBS, 100 U/mL penicillin/streptomycin and with recombinant IL-2 (rIL-2, 100 U/mL). All cells were maintained at 37°C under 5% CO_2_.

### Plasmids and proviral constructs

The plasmid encoding the soluble CD4 domains 1 and 2 (sCD4^D1D2^) fused to a His6x-tag, as well as the molecular clones of the heavy and light chains of the anti-HIV- 1 Env monoclonal antibodies 17b and 2G12, were kindly provided by Dr. Peter Kwong (NIAID, NIH). Plasmids for expression of NESPylRS^AF^/hU6tRNA^Pyl^ and eRF1-E55D for the *amber* codon suppression system were previously described (34). The pNL4-3 Δ*RT* Δ*env* plasmid has been previously described (11). pNL4-3.Luc.*R*-*E*- provirus was obtained from the former NIH AIDS Reagent Program (catalog ARP-3418). The stop codon in *tat* gene of this plasmid was substituted with an *ochre* stop codon as described

(39). Plasmids for the expression of full-length HIV-1_JR-FL_ Env wild-type, which was engineered to have an amber (TAG) stop codon at position N135 in the V1 loop of gp120 and the A1 peptide (GDSLDMLEWSLM) in the V4 loop of gp120 (V1- N135^TAG^/V4-A1) have been previously described (30). The HIV-1 _CRF01_AE_ Env expressor has been described (12). This plasmid was engineered to insert the A4 peptide (DSLDMLEW) after residue V135 in the V1 loop of gp120, and substitute an amber codon at position N398 in the V4 loop of gp120 (A4-V1/V4-N398^TAG^, Fig. 2B). The H375S mutation, was introduced into the untagged and tagged versions of the HIV- 1_CRF01_AE_ Env expression plasmids. All the indicated residues in HIV-1_JR-FL_ and HIV- 1_CRF01_AE_ Env are numbered according to the HIV-1_HXBc2_ Env sequence.

The infectious molecular clone (IMC) of HIV-1_JR-FL_ was kindly provided by Dr Dennis Burton (The Scripps Research Institute). The CRF01_AE IMC was previously reported (doi: 10.1128/JVI.02452-16). The sequence of HIV-1_CRF01_AE_ transmitted- founder (T/F) clone 703357 was derived by using a single-genome amplification (SGA) strategy. The entire DNA sequence including both long terminal repeats (LTRs) was cloned into pUC57 to generate a full-length infectious molecular clone (GenBank accession numbers JX448154 and JX448164). The vesicular stomatitis virus G (VSV-G)-encoding plasmid was previously described (46).

### Recombinant sCD4^D1D2^ and antibodies

Expression of soluble CD4 domains D1-D2 (sCD4^D1D2^) fused to a His6x-tag was performed by transfection of ExpiCHO-S™ cells with plasmid using the ExpiFectamine™ CHO transfection kit (Gibco™, Thermo Fisher Scientific, Waltham, MA, USA) according to the manufacturer’s instructions. Purification and preparation of this protein was performed with a previously described strategy(40). Briefly, supernatant containing soluble sCD4^D1D2^ was harvested nine days post-transfection and adjusted to 1 mM NiSO_4_, 20 mM imidazole, and pH 8.0 before binding to the Ni-NTA resin (Invitrogen™, Waltham, MA, USA). The resin was washed, and sCD4^D1D2^ was eluted from the column with 300 mM imidazole, 500 mM NaCl, 20 mM Tris-HCl pH 8.0, and 10% (v/v) glycerol. Elution fractions containing sCD4^D1D2^ were pooled and concentrated by centrifugal concentrators (Sartorius AG, Göttingen, Germany). Final purification was performed through size exclusion chromatography on a Superdex 200 Increase 10/300 GL column (GE Healthcare, Chicago, IL, USA) followed by concentration as above described.

Expression and preparation of monoclonal antibodies 2G12 and 17b has been described before (40, 41). Briefly, ExpiCHO-S cells were co-transfected with plasmids encoding heavy and light chains using the ExpiFectamine CHO transfection kit (Gibco, ThermoFisher Scientific, Waltham, MA, USA) according to the manufacturer’s instructions. Both antibodies were purified from the cell culture supernatant 12 days post-transfection using protein G affinity resin (Thermo Fisher Scientific, Waltham, MA, USA), subjected to buffer exchange with phosphate buffered saline (PBS) pH 7.4 (Fisher Bioreagents, Thermo Fisher Scientific, Waltham, MA, USA) and concentrated as above described. Mouse monoclonal antibody targeting HIV-1 p24 capsid protein (anti- p24, catalog No. GTX41618) was purchased from Genetex (Irvine, CA, USA). Anti-6x- His-tag polyclonal antibody (catalog No. PA1-983B), horseradish peroxidase (HRP) conjugated anti-human IgG Fc (catalog No. A18823), and anti-mouse IgG Fc (catalog No. 31455) were purchased from Invitrogen™ (Waltham, MA, USA). Goat anti-rabbit IgG antibody conjugated to HRP (catalog No. ab205718) were purchased from Abcam (Cambridge, UK).

### Virus production and fluorescent labeling

Non-replicative HIV-1_CRF01_AE_ Env pseudoviruses for infectivity assays were produced by co-transfecting HEK293T-FIRB cells with either a 1:0.005 or 1:1 mass ratio of plasmid pNL4-3.Luc.R-E- *tat*_*ochre* to wild-type or V1-A4/V4-N398^TAG^ tagged version of HIV-1_CRF01_AE_ Env expressors, respectively. Plasmids encoding NESPylRS^AF^/hU6tRNA^Pyl^ and eRF1-E55D were also included along with 0.5 mM TCO* (SiChem GmbH, Bremen, Germany) as previous described (30, 39, 42, 43). Virus was collected 48 hours post-transfection and pelleted over a 10% sucrose cushion at 25,000 RPM for 2 hours at 4 °C using a SW32Ti rotor (Beckman Coulter Life Sciences, Brea, CA, USA). Pellets were resuspended in DMEM (Gibco ThermoFisher Scientific, Waltham, MA, USA), aliquoted, and stored at -80 °C until use.

For smFRET imaging, non-replicative HIV-1_JR-FL_ and HIV-1_CRF01_AE_ Env pseudovirions with a single gp120 domain bearing the above-mentioned modifications in the V1 and V4 loops, were also produced in the presence of TCO* as previously described (30). Briefly, HEK-293T FIRB cells were co-transfected with plasmids NESPylRS^AF^/hU6tRNA^Pyl^ and eRF1-E55D, in addition of pNL4-3 ΔRT ΔEnv, and a 20:1 mass ratio of HIV-1_JR-FL_ or HIV-1_CRF01_AE_ Env wild-type expressor to the corresponding tagged version. Virus was collected 48 hours post-transfection and pelleted as above. Virus pellets was then resuspended in labeling buffer (50 mM HEPES pH 7.0, 10 mM CaCl_2_, 10 mM MgCl_2_), and incubated overnight at room temperature with 5 μM LD650- coenzyme A (Lumidyne Technologies, New York,NY, USA), and 5 μM acyl carrier protein synthase (AcpS), which labels the A1 (or A4) peptide. Virus was then incubated with 0.5 µM Cy3-tetrazine (Jena Biosciences, Jena, Germany) for 30 min at room temperature, followed by incubation with 60 μM DSPE-PEG2000-biotin (Avanti Polar Lipids, Alabaster, AL, USA) for an additional 30 min at room temperature. Finally, labelled virus was purified through ultracentrifugation for 1 hour at 35,000 RPM using a rotor SW40Ti (Beckman Coulter Life Sciences, Brea, CA, USA), at 4 °C in a 6–30% OptiPrep (Sigma-Aldrich, MilliporeSigma, Burlington, MA, USA) density gradient. Labelled pseudovirions were collected, analyzed by anti-p24 Western blot, aliquoted, and stored at -80°C until their use in imaging experiments.

### Immunoblots

HIV-1 gp120 and p24 proteins, or sCD4^D1D2^ were detected through immunoblot assays as follows. Samples were mixed with 4X Laemmli sample buffer (Bio-Rad, Hercules, CA, USA) supplemented with 2-mercaptoethanol (Fisher Chemical, Hampton, NH, USA) and heated for 5 min at 98 °C. Proteins were then resolved by denaturing PAGE using 4-20% acrylamide gels (Bio-Rad, Hercules, CA, USA). Proteins were then transferred to nitrocellulose membranes (Bio-Rad, Hercules, CA, USA) according to the manufacturer’s instructions. After blocking for one hour at room temperature with 5% (w/v) skim milk in PBS-T buffer [PBS and 0.1% (v/v) Tween™-20, Fisher Scientific, Hampton, NH, USA], membranes were incubated overnight at 4 °C with the indicated primary antibodies diluted in blocking buffer. Detection of gp120 was achieved by using a 3 µg/ml dilution of 2G12, while detection of p24 and sCD4^D1D2^ was performed with 2 µg/ml dilutions of anti-p24 mAb (GeneTex, Irvine, CA, USA) or rabbit anti-6x-His-tag polyclonal antibody (Invitrogen™, Waltham, MA, USA), respectively. Membranes were washed three times with PBS-T and incubated for one hour at room temperature with a 1/10,000 dilution (v/v) in 0.5% (w/v) skim milk/PBS-T of HRP-conjugated anti-human IgG Fc or anti-mouse IgG Fc (Invitrogen™, Waltham, MA, USA) antibodies for membranes incubated with 2G12 or anti-p24 mAbs, respectively, or a 1/50,000 dilution of HRP-conjugated anti-rabbit IgG antibody(Abcam, Cambridge, UK) was used for membranes incubated with anti-His6X antibody. After three washes with PBS-T, membranes were developed using SuperSignal™ West Pico PLUS Chemiluminescent Substrate (Thermo Scientific™, Waltham, MA, USA) according to the manufacturer’s instructions.

### Infectivity assays

2.5x10^4^ TZM-bl cells/well were seeded 24 hours before the assay in 24-well plates. Cells were then washed once with DMEM (Gibco, ThermoFisher Scientific, Waltham, MA, USA) and inoculated with pseudo-typed lentiviruses bearing wild-type or tagged HIV-1_CRF01_AE_ Env. After 2 h of virus adsorption at 37 °C, viral inoculums were removed and cells were washed with DMEM, followed by addition of fresh complete phenol red-free DMEM (Gibco, ThermoFisher Scientific, Waltham, MA, USA). Cell supernatants were removed 48 hours post-infection. The cells were lysed with Glo Lysis Buffer (Promega, Madison, WI, USA) according to the manufacturer’s instructions. Luciferase activity in cell lysates was detected by mixing equal volumes of lysate and Steady-Glo Luciferase Assay System reagent (Promega, Madison, WI, USA) and measured on a Synergy H1 microplate reader (Biotek, Winooski, VT, USA). The luminescence signal from mock infected cell lysates was subtracted from the signal obtained from infected cells and normalized by the abundance of both envelope gp120 and p24 proteins in viral inoculums, which were determined through densitometric analysis of protein bands observed in immunoblots using ImageJ software v1.52q (NIH, Bethesda, MD, USA). Infectivity was expressed as the percentage of that seen in cells inoculated with wild-type HIV-1_CRF01_AE_ Env pseudovirions.

### smFRET Imaging

Labelled HIV-1_JR-FL_ or HIV-1_CRF01_AE_ Env pseudovirions were immobilized on streptavidin-coated quartz slides and imaged on a custom-built wide-field prism-based TIRF microscope (39, 44). Where indicated, pseudovirions were incubated with 50 µM sCD4^D1D2^ and 50 µg/ml 17b mAb for 1 hour at room temperature prior to surface immobilization. Imaging was performed in phosphate-buffered saline (PBS) pH ∼7.4, containing 1 mM trolox (Sigma-Aldrich, St. Louis, MO, USA), 1 mM cyclooctatetraene (COT; Sigma-Aldrich, St. Louis, MO, USA), 1 mM 4-nitrobenzyl alcohol (NBA; Sigma-Aldrich, St. Louis, MO, USA), 2 mM protocatechuic acid (PCA; Sigma-Aldrich, St. Louis, MO, USA), and 8 nM protocatechuate 3,4-deoxygenase (PCD; Sigma-Aldrich, St. Louis, MO, USA) to stabilize fluorescence and remove molecular oxygen. When indicated, concentrations of sCD4^D1D2^ and mAb 17b were maintained during imaging. smFRET data were collected using Micromanager v2.0 at 25 frames/sec, processed, and analyzed using SPARTAN software in Matlab (Mathworks, Natick, MA, USA) (45). smFRET traces were identified according to criteria previously described (8); traces meeting those criteria were verified manually. FRET histograms were generated by compiling traces from each of three technical replicates and the mean probability per histogram bin ± standard error was calculated. Traces were idealized to a five-state HMM (four nonzero-FRET states and a zero-FRET state) using the maximum point likelihood (MPL) algorithm (46). The idealizations were used to determine the occupancies (fraction of time until photobleaching) in each FRET state, and construct Gaussian distributions of each FRET state, which were overlaid on the FRET histograms to visualize the results of the HMM analysis. The distributions in occupancies were used to construct violin plots in Matlab, as well as calculation of mean occupancies and standard errors.

### Viral production and infection of primary CD4+ T cells

VSV-G-pseudotyped HIV-1 viruses were produced by co-transfection of 293T cells with the HIV-1_JRFL_ or HIV-1_CR01AE_ proviral construct and a VSV-G-encoding vector at a ratio of 3:2 using the polyethylenimine (PEI) method. Two days post-transfection, cell supernatants were harvested, clarified by low-speed centrifugation (300 × g for 5 min), and concentrated by ultracentrifugation at 4°C (100,605 × g for 1h) over a 20% sucrose cushion. Pellets were resuspended in fresh RPMI 1640 complete medium, aliquoted and stored at -80°C until use.

Primary CD4+ T cells from HIV-1 negative individuals were isolated from PBMCs, activated for 2 days with PHA-L and then maintained in RPMI 1640 complete medium supplemented with rIL-2. Five to seven days after activation, the cells were spinoculated with the virus at 800 × g for 1h in 96-well plates at 25°C. All viral productions were titrated on primary CD4+ T cells to achieve similar levels of infection (around 20% of infected cells).

### Flow cytometry analysis of cell-surface staining

Forty-eight hours after infection, HIV-1-infected primary CD4+ T cells were collected, washed with PBS and transferred in 96-well V-bottom plates. The cells were then incubated for 45 min at 37°C with plasma (1:1000 dilution. Cells were then washed twice with PBS and stained with anti-human IgG Alexa Fluor 647-conjugated secondary antibody (2 μg/mL), FITC-conjugated mouse anti-human CD4 (Clone OKT4) Antibody (1:500 dilution) and AquaVivid viability dye (Thermo Fisher Scientific, Cat# L43957) for 20 min at room temperature. Alexa-Fluor-conjugated anti-human IgG Fc secondary antibodies (1:1500 dilution) were used as secondary antibodies. Cells were then washed twice with PBS and fixed in a 2% PBS-formaldehyde solution. The cells were then permeabilized using the Cytofix/Cytoperm Fixation/Permeabilization Kit (BD Biosciences, Mississauga, ON, Canada) and stained intracellularly using PE-conjugated mouse anti-p24 mAb (clone KC57; Beckman Coulter, Brea, CA, USA; 1:100 dilution).

Samples were acquired on an Fortessa cytometer (BD Biosciences), and data analysis was performed using FlowJo v10.5.3 (Tree Star, Ashland, OR, USA). The percentage of productively infected cells (p24^+,^ CD4^-^) was determined by gating on the living cell population according to viability dye staining (Aqua Vivid; Thermo Fisher Scientific).

### ADCC assay

ADCC activity was measured using a FACS-based infected cell elimination assay 48 hours after infection. The HIV-1-infected primary CD4+ T cells were stained with AquaVivid viability dye and cell proliferation dye eFluor670 (Thermo Fisher Scientific) and used as target cells. Resting autologous PBMCs, were stained with cell proliferation dye eFluor450 (Thermo Fisher Scientific) and used as effectors cells. The HIV-1- infected primary CD4+ T cells were co-cultured with autologous PBMCs (Effector: Target ratio of 10:1) in 96-well V-bottom plates in the presence of plasma from PLWH (dilution 1:1000) for 5h at 37°C. After the 5h incubation, cells were then washed once with PBS and stained with FITC-conjugated mouse anti-human CD4 (Clone OKT4) antibody for 10 min at room temperature. Cells were then washed twice with PBS and fixed in a 2% PBS-formaldehyde solution. The cells were then permeabilized and stained intracellularly for p24 as described above. Samples were acquired on a Fortessa cytometer (BD Biosciences), and data analysis was performed using FlowJo v10.5.3 (Tree Star, Ashland, OR, USA). The percentage of infected cells (p24^+,^ CD4^-^) was determined by gating on the living cell population according to viability dye staining (Aqua Vivid; Thermo Fisher Scientific). The percentage of ADCC was calculated with the following formula: [(% of p24^+^CD4^-^ cells in Targets plus Effectors) − (% of p24^+^CD4^-^ cells in Targets plus Effectors plus plasma)/(% of p24^+^CD4^-^ cells in Targets) × 100].

### Statistical analysis

Statistics for infectivity assays were determined using GraphPad Prism version 10.2.3 (GraphPad, San Diego, CA, USA). Every data set was tested for statistical normality and this information was used to apply the appropriate (parametric or nonparametric) statistical test. Statistical significance measures (*p*-values) of FRET state occupancies were determined by one-way ANOVA followed by multiple comparison testing in Matlab (The MathWorks, Waltham, MA, USA). In all cases, *p*- values <0.05 were considered statistically significant.

## Acknowledgements

The authors thank Dr. Robert Blakemore for assistance in the design of the tagged HIV-1_CRF01_AE_ Env glycoprotein, Dr. Dennis Burton (The Scripps Research Institute) for HIV-1_JRFL_, and Dr. Peter Kwong (NIAID, NIH) for providing the molecular clones of sCD4^D1D2^, and mAbs 2G12 and 17b heavy and light chains. The TZM-bl cell line (ARP-8129) was obtained through the former NIH HIV Reagent Program, Division of AIDS, NIAID, NIH, and this cell line was originally contributed to them by Dr. John C. Kappes, Dr. Xiaoyun Wu and Tranzyme Inc. M.N. was supported by a ViiV postdoctoral fellowship. J.P. was supported by a CIHR fellowship. This work was supported by NIH grant R01AI150322 (to J.B.M. and A.F.).

## Author Contributions

J.B.M. and A.F. conceived of the study. M.A.D.-S., D.C., M.N., H.M., J.P., M.P., A.F., and J.B.M. designed experimental approaches, performed, analyzed, and interpreted the experiments. A.F., and J.B.M. obtained the funding and supervised the research. M.A.D.-S., and J.B.M. wrote the manuscript. All authors have read, edited, and approved the final manuscript.

## Disclaimer

The views expressed in this manuscript are those of the authors and do not reflect the official policy or position of the Uniformed Services University, US Army, the Department of Defense, or the US Government. The funders had no role in study design, data collection and analysis, decision to publish, or preparation of the manuscript.

## Data Availability

All data generated or analyzed during this study are included in the manuscript.

